# Deep structural analysis of rhamnogalacturonan-I (RG-I) in Arabidopsis reveals new insights into pectin diversity

**DOI:** 10.1101/2024.10.25.620358

**Authors:** Liang Zhang, Jiri Vlach, Ian M. Black, Stephanie Archer-Hartmann, Christian Heiss, Parastoo Azadi, Breeanna R. Urbanowicz

**Affiliations:** Complex Carbohydrate Research Center, University of Georgia, Athens, GA 30602, USA; Department of Biochemistry and Molecular Biology, University of Georgia, Athens, GA 30602, USA

## Abstract

Pectic polysaccharides are an integral part of primary plant cell walls, where they are perfrom crucial structural and biological functions. Pectin is generally divided into four distinct structural categories, including homogalacturonan, xylogalacturonan, rhamnogalacturonan I (RG-I) and rhamnogalacturonan II. Among the four main pectin domains, the structural intricacies of rhamnogalacturonan-I (RG-I) remain the least understood, especially in widely employed plant models. We employed multiple complementary analytical techniques to present a detailed structural analysis of RG-I in the model system *Arabidopsis thaliana.* Using highly purified RG-I from different tissues, we conducted comparative linkage and NMR analyses, complemented by mass spectrometry of enzymatically digested RG-I oligosaccharides. Our findings present the most comprehensive structural overview of Arabidopsis RG-I to date, revealing novel structural features. Notably, we identified *O*-acetylation of rhamnose backbone residues as a predominant feature, a modification previously unreported in this species. The combined results present a comprehensive structural overview of *Arabidopsis thaliana* RG-I that will serve as a roadmap for studying pectin biosynthesis and function.

## Introduction

All plant cells are surrounded by a cell wall, mainly composed of an intricate network of polysaccharides that are not only structural elements but also key players in plant physiology and adaptation. Plants are multicellular organisms that rely on their cell walls to play many different roles, including facilitating growth and cell-cell interactions, providing mechanical support and dynamic extensibility, protecting against pathogens, and regulating water movement (Cosgrove 2024). Pectins are a group of plant cell wall polysaccharides that represent the main components in the continuous middle lamella layer present between neighboring cells, and are often highly abundant in primary walls and seed mucilage of dicots and nongraminaceous monocots (Jarvis et al. 2003; Ferrari et al. 2013; Feng et al. 2018). Lower amounts of pectins are also found in secondary cell walls and in the primary walls of grasses. In current cell wall models, it is proposed that pectins form a gel-like matrix, embedding cellulose microfibrils and hemicellulose (Lampugnani et al. 2018; Cosgrove 2024). Evidence has shown that pectins covalently or noncovalently interact with cellulose, hemicellulose, and arabinogalactan proteins (AGPs) (Thompson and Fry 2000; Abdel-Massih et al. 2003; Oechslin et al. 2003; Tan et al. 2013; Lin et al. 2016; Broxterman and Schols 2018; Tan et al. 2023), indicating these polysaccharides play important roles in cell wall assembly and the maintenance of its integrity. Plant cell activities such as growth, differentiation, and response to environmental signals require the plant cell wall to be extensible. The high water-holding capacity of pectins and their mild interactions with other cell wall polysaccharides provide the wall with such extensibility (Cosgrove 2022); therefore, pectins are important in a wide variety of biological contexts. Mutant plants that produce altered pectins exhibit developmental defects, including, but not limited to, reduced size of rosette leaves (O’Neill et al. 2001; Rui et al. 2017), disrupted organ initiation and asymmetric cell growth (Peaucelle et al. 2011; Bou Daher et al. 2018), and impaired pollen tube elongation (Jiang et al. 2005).

A key reason why we generally refer to ‘pectins’ as so complex is that they are defined by the fine structures and abundance of their four pectin domains: homogalacturonan (HG), xylogalacturonan (XG), rhamnogalacturonan I (RG-I) and rhamnogalacturonan II (RG-II). RG-I is unique among pectins as its backbone is formed by a disaccharide repeat of 2-α-L-rhamnose (Rha) and 4-α-D-galacturonic acid (GalA), →2)-α-L-Rha(1→4)-α-D-GalA(1→. On the basis of current generalized models of RG-I, Rha residues can be substituted at *O*-4 with linear or branched arabinan, galactan, or arabinogalactan side chains. The arabinan side chains have a linear core of 5-linked α-L-arabinofuranose (Ara*f*) residues that can be decorated at *O*-2 or/ and *O*-3 with single α-L-Ara*f* residues or with short chains of 3-linked α-L-Ara*f* residues (Yapo 2011). Galactan side chains are usually made of 4-linked β-D-galactose (Gal) residues (Yapo 2011). However, Gal residues can be branched at *O*-3 with single α-L-Ara*f* residues or with short chains of 5-α-L-Ara*f* residues, thus forming type I arabinogalactan (Yapo 2011). Type II arabinogalactans differ from type I arabinogalactans in two ways. First, the backbone consists of 3-linked β-D-Gal residues. Second, type II arabinogalactans are more branched than type I arabinogalactans. Gal residues in type II arabinogalactans are substituted at *O*-6 with single Gal residues or with short chains of 6-linked β-D-Gal residues, which can be further substituted at *O*-3, *O*-4 or/ and *O*-6 with α-L-Ara*f* containing chains (Yapo 2011). D-Glucuronic acid (β-D-GlcA) or 4-*O*-methyl-GlcA (4-*O*-Me-β-D-GlcA) residues that are 6-linked to Gal residues in type II arabinogalactans are also found (Tsumuraya et al. 1990). Substitution of GalA residues in RG-I is rare. A study on RG-I oligosaccharides made from sugar beet pulp showed that the backbone GalA residues were appended with β-D-GlcA at *O*-3 (Renard et al. 1999). RG-I is known to be modified by *O*-acetyl substituents that have been thought to predominantly occur at *O*-2 or/ and *O*-3 of the GalA residues (Ishii 1997), similar to HG. Acetylation at *O*-3 of Rha residues has also been reported in okra (Sengkhamparn et al. 2009), although it was believed to be unusual.

Arabidopsis has been used as a model plant system to study the structure, biosynthesis, remodeling, and function of the cell wall and its polysaccharides for more than 30 years (Reiter et al. 1997; Liepman et al. 2010). Despite an abundance of research on the biological role of pectins, our understanding regarding the fine structure of pectins is quite incomplete in this model species. Indeed, because of the combination of their complexity and functional importance for cell-cell adhesion and plant growth, the identification of mutants with altered pectin structure has lagged behind that of other wall polymers. Most *in-vivo* studies of pectins have focused on HG. However, the function of RG-I in plants remains unclear. It is widely believed that RG-I structure is diverse and varies by tissue, species, and developmental stages (Kaczmarska et al. 2022). This raises the intriguing question regarding what is the true complexity and diversity of RG-I in Arabidopsis, and does it reflect and even dictate its functions. Large gene families proposed to be involved in RG-I biosynthesis suggest that its biosynthesis likely involves enzyme homologs that are functionally redundant while exhibiting distinct characteristics (Harholt et al. 2012; Liwanag et al. 2012; Takenaka et al. 2018; Delmer et al. 2024). Our knowledge regarding RG-I structure has been compiled from non-model plant species, while very little is known for model species, creating challenges in identifying and studying enzymes that synthesize RG-I. Here, we performed deep structural analyses of RG-I isolated from different Arabidopsis tissues. We found that RG-I from this model plant was substituted with arabinan, galactan, and arabinogalactan side chains. In addition, we showed that GalA and Rha residues of the backbone were *O*-acetylated, with Rha substituted at *O*-3 being predominant in all tissues. Characterization of enzyme functions is often limited by substrate availability. We used an RG-I lyase to create small RG-I fragments for our mass spectrometry analysis. Those fragments can be used as substrates for studying the functions of RG-I biosynthetic enzymes.

## Results

### Arabidopsis RG-I is highly complex and contains linear and branched galactan, arabinan, and arabinogalactan

Polysaccharides are functional macromolecules with structures defined on the basis of the glycosyl sequence of their constituent monosaccharides. To define the key structural motifs of RG-I in Arabidopsis, cell walls were prepared as alcohol-insoluble residues (AIR) and used as the basis to develop optimized fractionation and analytical workflows for structural analysis of pectic fragments (Figure 1). To optimize our purification scheme, we took advantage of an optimized method for uronic acid linkage analysis that incorporates a preacetylation step in ionic liquid prior to the preparation of partially methylated alditol acetate (PMAA) derivatives (Black et al. 2023). We applied this method here, as traditional methods used to analyze RG-I were determined to be inaccurate because acidic polysaccharides are poorly soluble in DMSO and tend to undergo β-elimination during permethylation. To obtain a global representation of pectins in each tissue, AIR was treated with ammonium oxalate, and the pectin-enriched fraction was subjected to glycosyl linkage analysis with preacetylation (Black et al. 2023).

**Figure 1.**
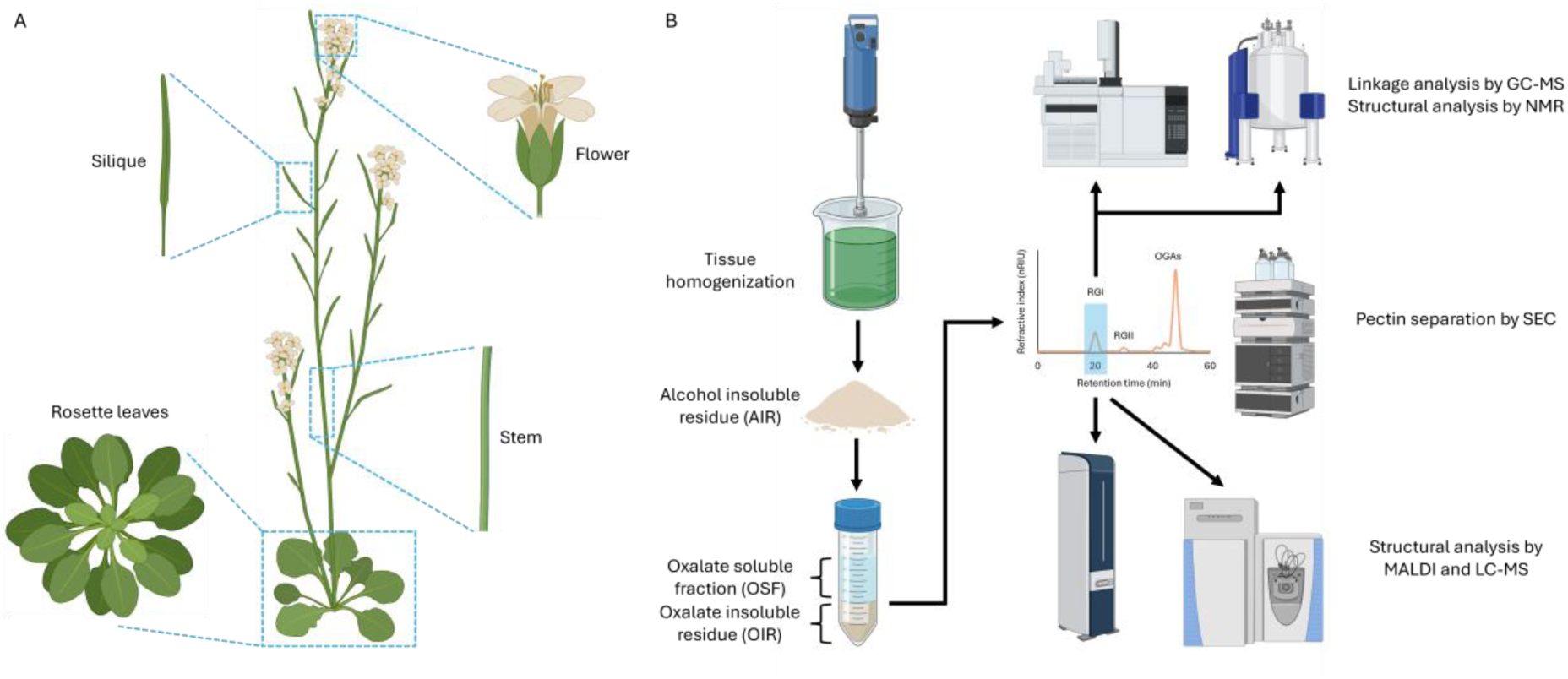
Pectin analysis workflow. **(A)** Schematic representation of Arabidopsis tissues used in this study. **(B)** General workflow for isolating pectin fragments from Arabidopsis cell walls, prepared as alcohol insoluble residues (AIR), for structural analyses. Pectin fragments were generated by a combination of chemical and enzymatic treatments prior to separation and purification by size exclusion chromatography (SEC). Fractions enriched in RG-I were subjected to structural analyses.

Our data showed that the oxalate soluble fractions (OSF) of leaf and silique AIR had about 3-4 times more 4-linked GalA, 2,4-linked GalA, and 3,4-linked GalA than 2-linked Rha and 2,4-linked Rha (Supplementary Figure 1), suggesting that the solubilized pectins of leaf and silique contained more HG than RG-I. NMR analysis of these same samples showed strong signals of HG across all four tissue types, whereas the signals of RG-I, which is represented by Rha, were weaker (Supplementary Figure 2). Next, we digested the soluble oxalate fractions using endo-polygalacturonanase (EPG) to release RG-I. Prior work in our lab found that after oxalate extraction, a significant amount of pectins remain present in the oxalate insoluble residue (OIR) (Barnes et al. 2021). Thus, oxalate insoluble residues (OIR) were treated with EPG, RG-I was separated by size exclusion chromatography (SEC), and collected (Supplementary Figure 3).

More RG-I was obtained from the oxalate insoluble residue (OIR) than from the oxalate soluble fraction (OSF) per unit of EPG (Supplementary Figure 3); thus, we decided to focus our studies on RG-I isolated from the oxalate insoluble residue (OIR), from here on out referred to as RG-I. The extraction capacity of RG-I and pectins in ammonium oxalate can be correlated with their structural properties, so it is important to consider both fractions when evaluating pectin structures from plant samples. Linkage analysis showed that the GalA:Rha ratio was close to 1 for RG-I generated from all four tissues (Supplementary Figure 1). We observed 5-linked Ara*f* and 4-linked Gal, indicating the presence of 5-linked α-arabinan and 4-linked β-galactan side chains. Furthermore, structures containing 2,5-linked Ara*f*, 3,5-linked Ara*f*, 2,3,5-linked Ara*f* and 3,4-linked Gal were identified, indicating that the arabinan and galactan side chains were branched. Since a larger amount of 5-linked Ara*f* and 4-linked Gal was present in these fractions relative to their multi-linked siblings, the linear forms were predominant. Type II arabinogalactan, which is characterized by 3-, 3,6-, and 6-linked Gal, was also identified.

We then analyzed RG-I obtained from OIR by NMR (Supplementary figure 4). Prior reports suggest that AGPs isolated from Arabidopsis cell culture and plant tissues are linked to RG-I (Tan et al. 2023; Tan et al. 2024); however, signals corresponding to proteins were absent from all NMR spectra (Supplementary figure 4). Indeed, AGPs and pectic-AGPs are typically solubilized during the enzymatic de-starching process prior to pectin isolation (Smith et al. 2020; Tan et al. 2024). Our data suggest that the optimized isolation protocol developed herein enriches RG-I that is not covalently linked to AGP. We observed signals characteristic of RG-I as well as GalA signals characteristic of acetylated and methyl-esterified HG, that were prominent likely due to the higher flexibility of HG regions and partially overlapped with the GalA signals from RG-I. To better assign RG-I peaks, we treated RG-I again with EPG but with the addition of pectin methylesterase (PME). PME removes methyl-esters from HG GalA residues, which may have hindered the activity of EPG (Esquivel and Voget 2004). The ^1^H and ^1^H,^13^C-HSQC NMR spectra of the resulting RG-I derived from the four different tissues showed that a second enzymatic digestion yielded RG-I free of most HG contamination, with the exception of leaf (Figure 2 B and C), which contained substantial amounts of highly acetylated, EPG resistant HG. The purity of the samples was determined by the GalA:Rha ratio, which should be 1 for RG-I, and approached this value in all samples (Supplementary Figure 1). In addition to the presence of residues characteristic for RG-I, as described above, the linkage data also showed several low-abundance residues consistent with the presence of RG-II, (xylo)galacturonan and hemicelluloses, indicating that trace amounts of these glycans were still present with the purified RG-I (Figure 2A); these components likely co-purified with RG-I in the high molecular weight SEC fraction.

**Figure 2.**
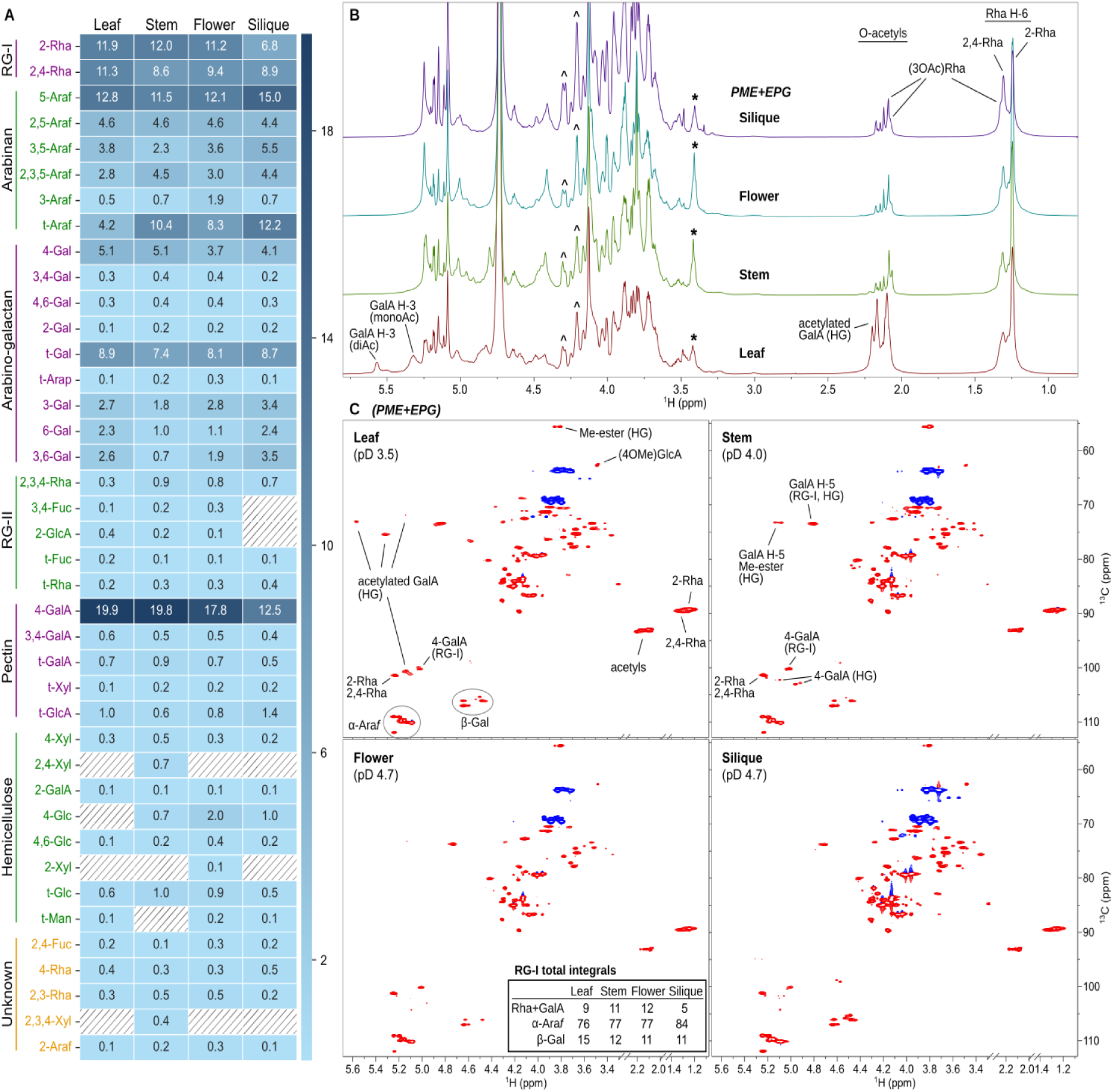
Analysis of RG-I from Arabidopsis tissues. **(A)** Glycosyl linkage analysis of RG-I (OIR 2). Colors refer to relative GC-MS peak area percentages, with dark blue indicating a high value and light blue indicating a low value. Not detected is shown as diagonal lines. All tissue types contained residues that indicated the presence of arabinan and galactan RG-I sidechains. Presence of characteristic residues and linkages indicated that a small amount of RG-II and hemicellulose were still associated with RG-I after two rounds of separation. Additional residues that couldn’t be categorized as a particular polymer were also present. While the linkage analysis cannot distinguish 5-Ara*f* from 4-Ara*p*, the arabinans herein were reported as 5-Ara*f* based on NMR results. **(B)** A stack of ^1^H NMR spectra (800 MHz) of RG-I (OIR 2). Signals of acetylated Rha (*O*-acetyl and H-6 methyl groups) that are partially resolved in silique, flower and stem spectra, but less obvious in the leaf spectrum, are labeled. Key differences between the ^1^H spectra are due to different signal intensities, highlighting the higher proportion of unbranched Rha in flower and stem (*), and higher overall arabinan content in silique RG-I (^). **(C)** ^1^H,^13^C-HSQC NMR spectra of the same RG-I samples shown in panel B, with characteristic signal groups labeled in the leaf and stem spectra. The in-set table lists semi-quantitative HSQC signal integrals corresponding to the RG-I backbone (Rha+GalA) and side-chains (total Ara and total Gal).

Silique RG-I contained the highest proportion of side chains, as determined from the relative intensities of the 2-Rha and 2,4-Rha signals (Figure 2 A and B); thus, it was selected for detailed NMR characterization. To improve the sensitivity and resolution of NMR signals, data were acquired on an 800 MHz instrument at two different pD values (4.7 and 1.7) to help resolve pH-sensitive residues. Analysis of NMR spectra revealed the presence of at least 25 distinct sugar residues, most of which could be identified and linked based on their chemical shifts and interaction patterns (Supplementary Table 1). The characteristic HSQC anomeric signals of the RG-I backbone were accompanied by signals corresponding to α-arabinan, 4-β-galactan, and 6-β-galactan side chains (Figure 3). Arabinan sidechains consisted of a 5-α-L-Ara*f* core structure, partially substituted with single Ara*f* residues in the 2-, 3- and 2,3-positions. No 3-linked arabinan was present. Very weak signals corresponding to terminal β-L-Ara*f* (Shakhmatov et al. 2019) were detectable in silique and flower tissues, but its connectivity could not be established. The 4-β-galactan side chains had an average degree of polymerization of 4 to 5 residues, estimated by the ratio of the integrals of the HSQC signals of the terminal and the in-chain groups. A large portion of the 6-β-galactan sidechains was appended at *O*-3 with single Ara residues and was terminated at the 6-position with 4-*O*-methyl β-GlcA. Similar termination of galactan or arabinogalactan (AG) chains with 4-*O*-Me-GlcA has been previously reported (Tsumuraya et al. 1990; Rakhmanberdyeva et al. 2019). Single-residue β-Gal substituents on RG-I backbone were also identified (Mikshina et al. 2012; Golovchenko et al. 2022). Broadened peak(s) in the β-Gal region (marked with asterisk in Figure 3) could not be assigned to particular structures because of weak and overlapping signals. These signals could be due to the t-Gal groups of the 6-galactan or other Gal residues. Also, we were not able to find NMR signals corresponding to a 3-β-galactan, even though 3-Gal was detected by the linkage analysis. Analysis of RG-I from all four Arabidopsis tissues under investigation revealed the same pattern of HSQC signals indicating that the structures of the arabinan, galactan and arabinogalactan side-chains were the same or very similar with differences only in relative amounts, but not structures of the substituents (Figure 2). All structural fragments identified by NMR are shown in Figure 3B, together with their chemical shift assignment codes (Supplementary Table 1).

**Figure 3.**
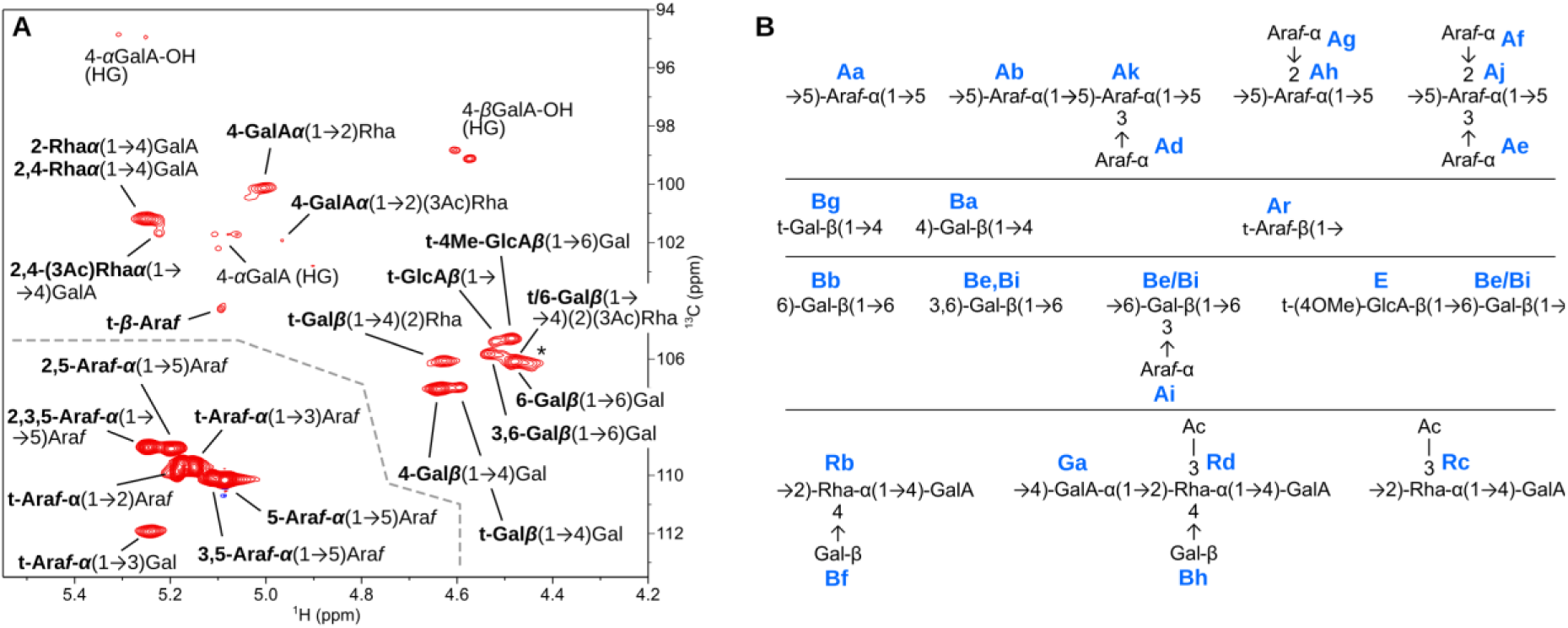
RG-I structural fragments identified in siliques. **(A)** Anomeric region of ^1^H,^13^C-HSQC spectrum of silique RG-I (OIR 2). The signal labels are shown as structural fragments where the residue highlighted in bold gives rise to the signal. Except for Ara, all residues are in the pyranose form. The dashed line serves as a visual aid to separate the arabinan signals from the rest. Asterisk marks broadened signals that could not be assigned to particular residues. **(B)** RG-I structural fragments identified by NMR. Each residue is labeled in blue with the code used in the chemical shift assignment table (Supplementary Table 1).

### 3-O-Acetylated rhamnose is the predominant O-acetyl modification of RG-I in Arabidopsis

In biological systems, molecules are often modified by additional functional groups during or after synthesis that modulate their chemical and functional properties. *O*-acetyl esterification is an important structural and functional feature of most plant and microbial polysaccharides and is found in the pectins present in cell walls of all terrestrial plants. This non-glycosyl substituents have been thought to occur at *O*-2 or *O*-3 of GalA residues in RG-I backbone. In contrast to the current dogma, examination of NMR spectra revealed signals due to 3-*O*-acetylation of Rha residues of the RG-I backbone (Figure 2 and Supplementary figure 5). TOCSY and NOESY spectra were especially instrumental in the assignments, despite lower intensity and overlaps of the (3*O*Ac)Rha signals (Supplementary Figure 5). Further, most of the 3-*O*-acetylated Rha backbone residues were substituted at the O-4 position with β-Gal substituents, though it was not possible to distinguish if they consisted of a single Gal or a chain of (1,6)-β-linked residues using NMR alone. A smaller proportion of (3*O*Ac)Rha was not branched. Abundant Rha acetylation was present in RG-I derived from all four tissues, as shown by comparative analyses of ^1^H spectra and 2D spectra (Figure 2), and to a similar degree among the tissues (7-11%) based on ^1^H integrals. Acetylation of Rha in RG-I was reported previously only for okra (Sengkhamparn et al. 2009), but our results suggest that such RG-I modification may be much more common, and is the predominant *O*-acetyl modification of all Arabidopsis RG-I analyzed herein. Our method of isolation of highly enriched RG-I as well as the use of high-resolution and high-sensitivity NMR instrumentation both enabled the detection of Rha acetylation which otherwise may have been missed. Besides the signals of acetylated Rha, there were several other, weaker acetyl group signals that could have originated from GalA in RG-I or HG, but we were not able to assign them to particular residues.

The NMR signals of backbone Rha and GalA showed broadening that could be attributed to the structural diversity of side chains and pH effects (signals sharpened at the lower pD). Besides the expected sensitivity of GalA signals to pD, the Rha H-5 signals also exhibited significant pD sensitivity (Supplementary table 1), possibly induced by neighboring GalA residues and a particular conformation of the RG-I backbone. In addition to the Rha H-5, slight pD sensitivity was observed also for the chemical shifts of the Rha acetyl groups. Through-space correlations (NOEs and ROEs) between the acetyl groups of Rha and ring atoms of the adjacent GalA revealed distinct chemical shifts of GalA residues affected by acetylation of the preceding (aglycon) Rha, and indicated a preferred conformation of the RG-I backbone. The observed inter-residue correlations were consistent with the helical secondary structural arrangement of RG-I backbone as was previously proposed (Makshakova et al. 2017; Zdunek et al. 2021).

### Enzymatic generation of RG-I fragments for structural studies

Decoding the structures of polysaccharides is often facilitated by chemical or enzymatic degradation of large, complex polymers into small oligosaccharides that are more amenable for analysis. Indeed, linkage specific enzymes are valuable tools that enable structural studies of complex pectins. To further investigate the structure of the arabinan side chains, we treated RG-I isolated from Arabidopsis tissues with endo-α-1,5-arabinanase (BT0368) (Luis et al. 2018), and showed that arabinose was released from RG-I isolated from all types of tissue except leaf. Interestingly, BT0368 was able to produce arabinose from leaf RG-I only after chemical de-esterification with alkali (Supplementary figure 6). Xylan and HG form helical structures (Walkinshaw and Arnott 1981b; Walkinshaw and Arnott 1981a; Busse-Wicher et al. 2014). It is possible that RG-I also form tertiary structures, which produce obstacles to RG-I degradation enzymes such as BT0368. Methyl-esterification at the *C*-6 carboxyl groups affects HG’s colloidal properties (Haas et al. 2020). The tertiary structures of RG-I may be affected by the acetyl groups on RG-I backbone and/or on the adjacent HG backbone, as leaf RG-I having the highest degree of acetylation on RG-I and the residual HG relative to the other tissues under investigation (Figure 2). The effect of HG acetylation on BT0368’s activity may be the consequence of a particular spatial distribution of HG relative to RG-I arabinan side chains and warrants further investigation. Treatment of Arabidopsis RG-I with endo-β-1,4-galactanase (BT4668) released galactobiose from all four tissue types (Supplementary figure 7). Unlike the activity of BT0368, BT4668 was able to cleave galactan from all RG-I samples without prior de-esterification. This may be due to functional difference between these two enzymes, or the influence of the substrate’s structure on enzyme activity.

To acquire more detailed information about the structure of RG-I, we sought to enzymatically deconstruct the polymer into small fragments for analysis by mass spectrometry (MS). To achieve this, we utilized an RG-I lyase (BT4175) (Luis et al. 2018), whose activity was verified for substituted and unsubstituted RG-I (Supplementary figure 8). BT4175 was active on both substrates, with higher activity on unbranched RG-I, which could be due to less steric hindrance or higher backbone content than in branched RG-I. We treated RG-I isolated from Arabidopsis flowers with BT4175. We first analyzed the raw digest by UPLC-ESI-MS. But the general signal intensity was low. Therefore, we enriched the products by anion exchange chromatography. Unfortunately, the ESI-MS signal intensity was not significantly improved. We noticed that RG-I fragments were usually detected in two forms in our ESI-MS experiments, one had ammonium adduct, another had sodium adduct (Supplementary figure 8), with the former as the dominant population. Our attempt to replace the ammonium adduct with sodium by spiking sodium chloride in our samples was not successful. Additionally, we noticed that ammonium adducts yielded more complicated MS^2^ fragmentation patterns than the corresponding sodium adducts; thus, obtaining MS^2^ data was challenging. Therefore, we performed MALDI-TOF MS analysis of these RG-I oligosaccharides. Signals were identified with m/z values characteristic of a DP2 RG-I oligomer substituted with galactan, arabinan or arabinogalactan side chains (Figure 4). In the m/z range of our analysis, galactan side chains had a DP of 1-10 hexose residues, and were directly appended to the backbone via Rha, based on the identification of daughter ions representing Rha-Gal, but not GalA-Gal, in MS^2^ spectra (Figure 5). The galactan side chains might be decorated with uronic acid, which in turn might be methyl etherified (Figures 4 and 5). According to our NMR data, we speculated that this uronic acid to be GlcA (Figure 3). Arabidopsis flower RG-I had peaks consistent with both arabinan and arabinogalactan side chains (Supplementary table 2). The largest arabinan side chain observed was DP7. Our MS analyses were unable to distinguish whether the arabinogalactan signals represented arabinans attached to the backbone via a single Gal residue, or arabinans with a terminal Gal residue, such as those found in soybean and potato RG-I (Nakamura et al. 2002; Øbro et al. 2004), due to low signal intensity of the parental ions.

**Figure 4.**
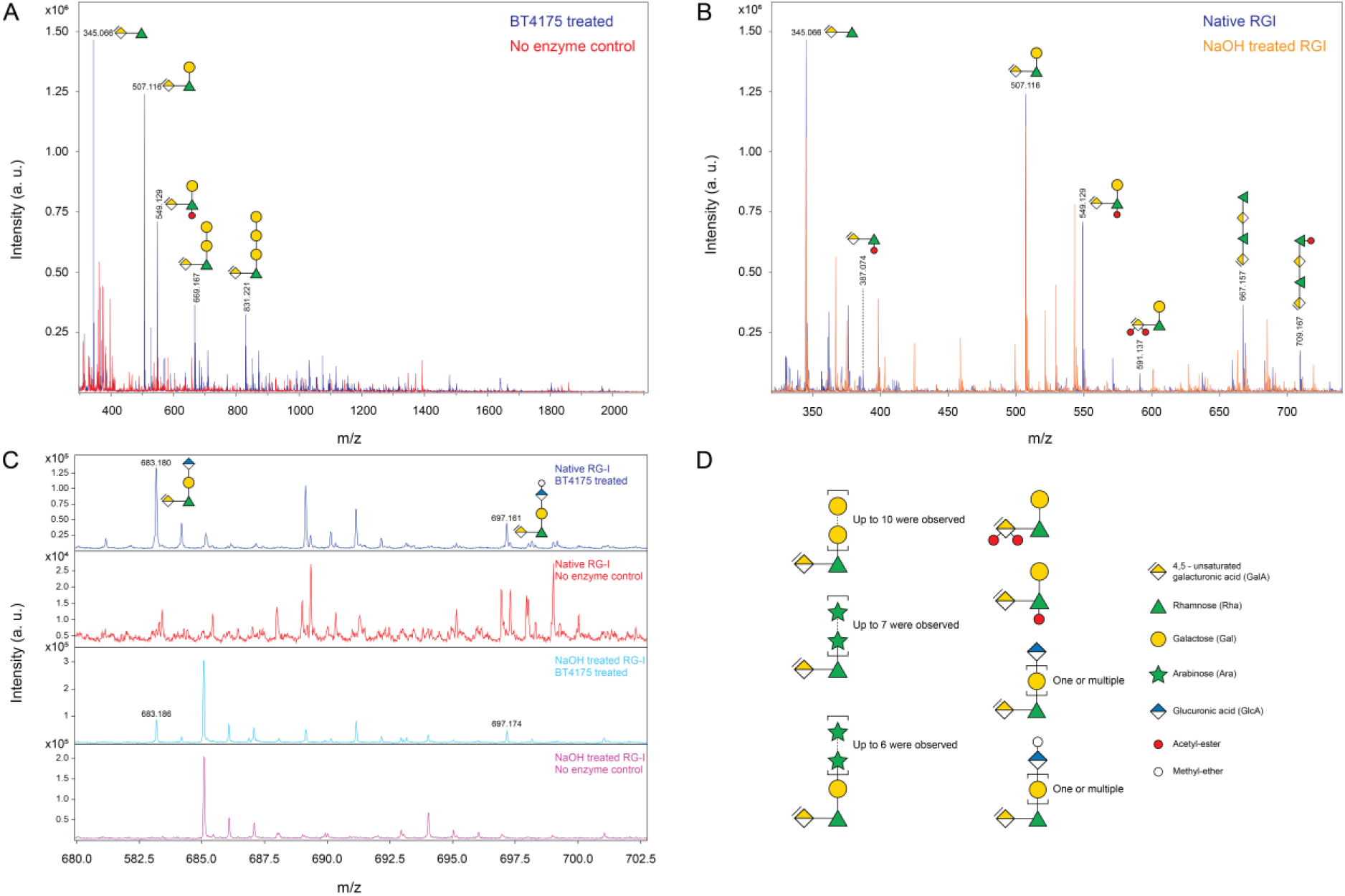
MALDI-TOF MS analysis of RG-I oligosaccharides from flower. **(A)** An overlay spectrum of BT4175 treated sample (blue) and the no enzyme control (red). Products made by BT4175 were not found in the no enzyme control. Only the most abundant peaks were labeled with their corresponding structures. **(B)** A representative spectrum showing oligosaccharides that had acetyl-substituents. These peaks were absent in the spectrum derived from the digestion of chemically de-esterified RG-I (orange). **(C)** A representative spectrum showing oligosaccharides that were substituted with side chains containing GlcA residues. These GlcA residue could be decorated with methyl-ether, since NaOH treatment was not able to eliminate it. **(D)** Schematic summarizing possible RG-I side chains and substituents present in flower tissue based on glycosyl linkage, NMR and MALDI-TOF MS analyses. More details about predicted and observed oligosaccharide masses can be found in Supplementary Table 2.

**Figure 5.**
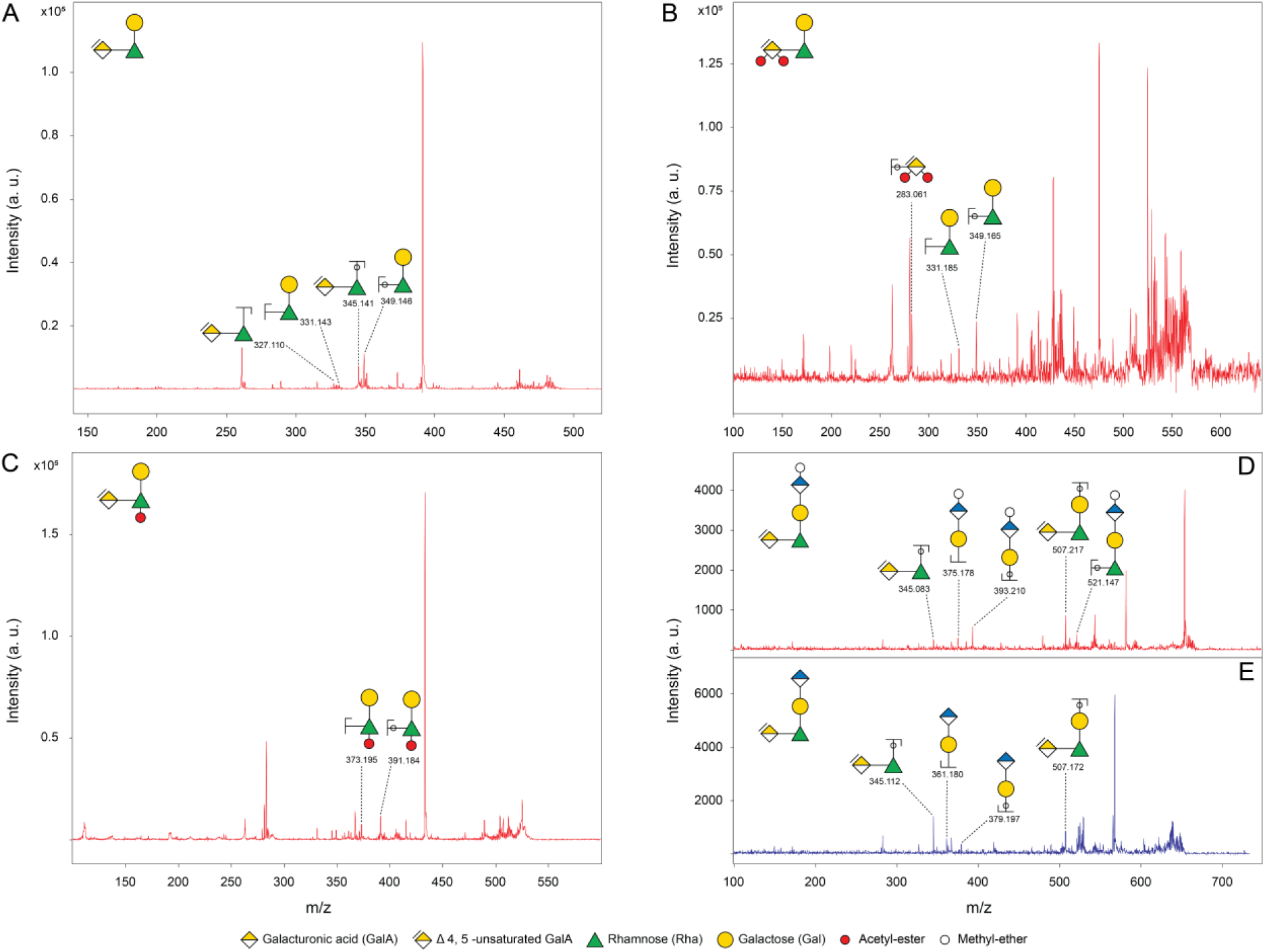
MALDI-TOF MS^2^ analysis of oligosaccharides produced from flower RG-I by RG-I lyase BT4175. **(A)** Galactose residue that was attached to Rha. **(B)** Doubly acetylated GalA at the non-reducing end. **(C)** Acetylation on Rha. **(D and E)** Side chains containing GlcA with (D) featuring a methyl-ether and (E) lacking this decoration.

Arabidopsis RG-I is believed to be further modified by *O*-acetyl substituents, which play a role in plant cell growth and interaction with the environment (Stranne et al. 2018). GalA residues *O*-acetylated at positions *O-*2 and/or *O-*3 were thought to be the major type of acetyl modification of the RG-I backbone, but most studies were carried out with non-model species (Perrone et al. 2002; Ralet et al. 2005). Indeed, a study on pectin structure in okra (*Abelmoschus esculentus*) identified Rha residues *O*-acetylated at position *O-*3 (Sengkhamparn et al. 2009); however, at the time it was believed to be uncommon. A key finding from this work is that Rha acetylation at *O-*3 is the dominant type of *O*-acetyl substituent in Arabidopsis RG-I (Figures 2 and 3). Our MS data show that the backbone of Arabidopsis flower RG-I contains *O*-acetyl groups on both Rha and GalA (Figure 5), and evidence of di-acetylated GalA was also given by MS^2^ (Figure 5).

This is surprising and indicates that there might be a multitude of functionally distinct RG-I *O*-acetyltransferases, including enzymes for GalA 2-*O*-acetylation, GalA 3-*O*-acetylation, Rha 3-*O*-acetylation, and potentially additional enzymes that can tolerate 4-linked Gal substituents on Rha and those involved in 2,3-*O*-acetylation of GalA. It is also likely that HG acetylation and RG-I acetylation require different sets of acetyltransferases. Structures identified by NMR showed that most Rha acetylation occurred on Rha residues that also contain glycosyl side chains at position *O*-4. Since acetylation can hinder the activity of enzymes that degrade plant cell wall polysaccharides (Gille and Pauly 2012), is it possible that Rha acetylation shields the branched regions in RG-I from being targeted by degradative enzymes? Or does this acetylation regulate substituent spacing and thus the local or global conformation of RG-I or the flexibility of the side chains, which in turn affects the interactions of RG-I with other cell wall components and the function of RG-I? Taken together, our data suggest that the BT4175 RG-I lyase tolerates both RG-I glycosyl side chains and backbone acetylation, thus representing a useful enzymatic tool to break down RG-I polysaccharides for structural studies.

### Hydrolytic enzymes can be used to build substrate library for studying RG-I biosynthesis

RG-I lyases catalyze β-elimination of RG-I polysaccharides, resulting in oligosaccharides containing an unsaturated GalA that absorbs UV light, which makes it convenient to monitor the enzymatic reactions and to separate the products. The activity of BT4175 relies on divalent cations (Supplementary figure 9). Incubation with EDTA led to loss of activity of BT4175 on Arabidopsis seed mucilage. The addition of Mg^2+^ or Ca^2+^ ion after the incubation recovered lyase activity, with Ca^2+^ ion showed greater effect, whereas the addition of Mn^2+^ ion made the lyase hyperactive. BT4175 belongs to the polysaccharide lyase family 11 (PL11). Other RG-I lyases in this family, such as Q65KY1_BACLD from *Bacillus licheniformis*, YesW and YesX from *Bacillus subtilis*, also show elevated activities in the presence of Mn^2+^ ions (Ochiai et al. 2007; Silva et al. 2011). BT4175 showed higher activity in alkaline conditions (Supplementary figure 9), which is consistent with other PL11 RG-I lyases (Ochiai et al. 2007; Silva et al. 2011).

But considering alkaline conditions might cause the loss of some structural information of RG-I, such as esterification, additionally, Mn^2+^ ion started to precipitate as Mn(OH)_2_ at pH 9, we decided to digest RG-I at neutral pH in the presence of Mn^2+^ ions. According to our MS data, the smallest product of BT4175 was RG-I disaccharide. It is important to obtain products having longer backbones, which can provide useful structural information about patterning of side chains and acetylation on the backbone. This can be achieved by reducing the activity of BT4175 with lower reaction temperature or lower concentration of Mn^2+^ (Supplementary figures 9 and 10). Since BT4175 tolerates both side chains and backbone acetylation, the RG-I oligosaccharides generated by this lyase from native RG-I polysaccharides can be used to build a substrate library for testing the activity of putative RG-I biosynthetic enzymes.

## Discussion

Due to the structural complexity of plant cell walls, estimations suggest 10% or more of the plant genome is dedicated to building the wall (Carpita and McCann 2015) Furthermore, genome-wide classification of carbohydrate-active enzymes (CAZymes; http://www.cazy.org) in Arabidopsis indicated more than 2% of the genome encodes large families of glycosyltransferases involved biosynthesis of complex carbohydrates (Drula et al. 2022). This make it difficult to address one of the most important research questions in plant biology: what are the structure-function relationships of cell wall polymers? This is an exceptionally difficult question to address, as the detailed structures of the majority of the saccharide components that comprise the cell wall remain undefined (Boerjan et al. 2024). In order to assign gene functions, baseline structural information is required to develop hypotheses and design experiments. We believe that a precise understanding of the glycosyl structures and sequences of complex polysaccharides, and clear approaches to study them, is required to move forward our understanding of their synthesis and biological importance.

Here we present a comprehensive structural model of the pectin RG-I present in the cell walls of Arabidopsis (Figure 6). Taken together, our combined data revealed the presence of several abundant structural features that were previously not known to exist in this important model plant species. Besides the presence of the canonical 5-arabinan, 4-galactan, 6-galactan and arabino-galactan RG-I side chains of varying lengths, we show that a large portion of the 6-galactan is terminated by either 4-*O*-methyl β-GlcA residues or, to a smaller degree, β-GlcA that lacks the Me-ether group. Importantly, we found that about 10% of the backbone Rha residues are 3-*O*-acetylated, while *O*-acetylation of RG-I GalA residues is less abundant. Most of the acetylated Rha is additionally substituted with β-Gal or galactan side chains. This comprehensive model was achieved using several complementary analytical techniques that compensate for methodological limitations inherent in any one analysis, allowing the elucidation of RG-I structural details with a high degree of certainty.

**Figure 6.**
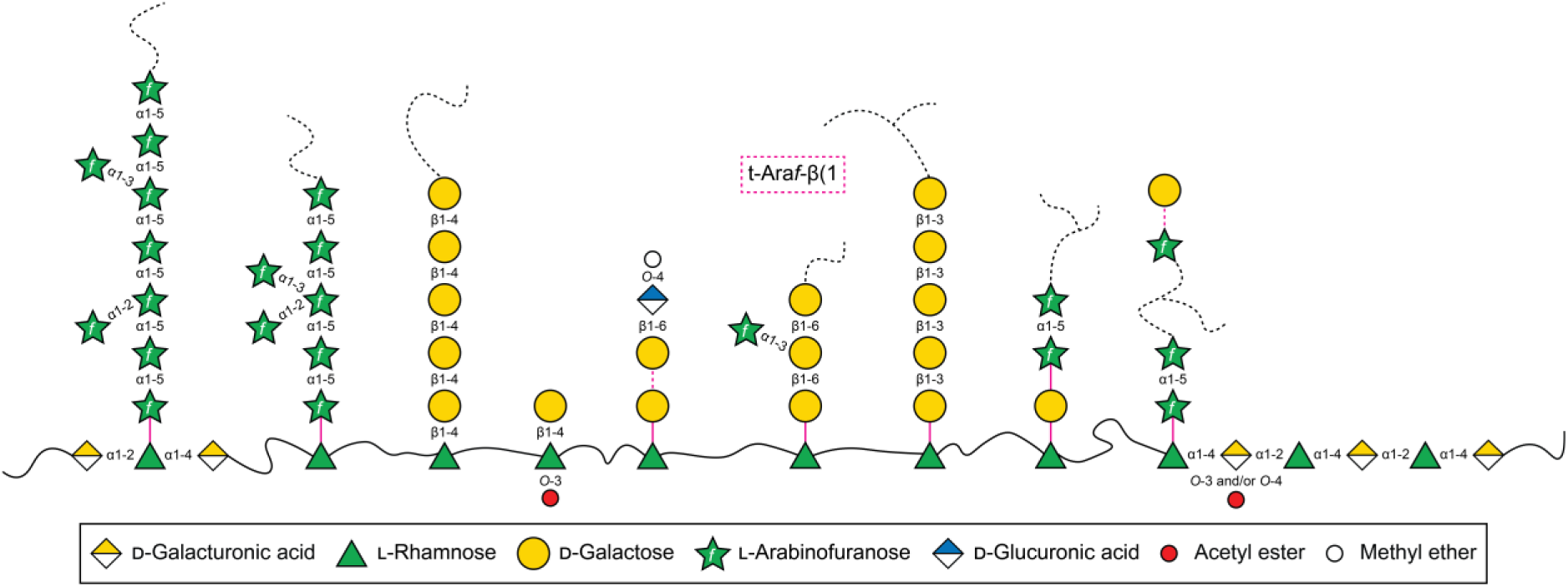
A schematic of RG-I structures in Arabidopsis. Arabinan side chains were substituted with single Ara*f* residues in 2-, 3-, and 2,3-positions. Arabinans were directly attached to the backbone, but the type of linkage is unknown. β1-4 and β1-6 linked galactan side chains were observed. β1-6 linked galactans were substituted with α1-3-linked Ara*f* residues. A portion of the galactan side chains were terminated by GlcA residues that carried methyl-ether at the *O*-4 position. Arabinogalactans were observed but it is not known how they are attached to the backbone. They may attach to the backbone via Gal or via Ara residue. The type of linkage to the RG-I backbone could be resolved by NMR only for the single-Gal side chain (β1-4) but not the other side chains. Acetylation on Rha of the backbone was the major acetyl modification of RG-I found in this study. Mono- and di-acetylated GalA residues were also observed. Terminal β-linked Ara*f* residue (in the magenta dash box) were observed, but where it was linked is unknown. Black line: simplified RG-I backbone. Black dash lines: the side chains can be longer and/or branched. Magenta lines: linkages not deciphered in this study. Magenta dash lines: linkages not deciphered, and the chains can have more than one Gal or Ara*f*.

However, we are aware that several key challenges still remain. One challenge in studying RG-I structure is the heterogeneity of the side chains, including their abundance, size, composition, and linkages. A question related to the heterogeneity of RG-I is what biological meaning it conveys? For example, why do some galactan side chains, but not all, have a terminal GlcA residue? Does this terminal GlcA affect the local hydration properties, local cell wall pH, or local availability of calcium ions? RG-I is structurally complex due to its degree and type of branching, as shown by previous studies and our results. We are unable to distinguish between the intramolecular and intermolecular complexities. According to our data, RG-I side chains are not uniform in size. For example, we observed galactan side chains varying in size from DP1 to DP10. Are the side chains of different sizes evenly distributed, or do they tend to cluster together with similar-sized side chains? Such questions regarding side chain patterning requires producing RG-I fragments with longer backbones. Another question is whether the heterogeneity in side chain size is biologically important. Do different sizes convey unique functions, or do they serve the same or similar functions but at varying levels? The heterogeneity in side chain size may be one reason why RG-I provides extensibility to the cell wall. Moreover, are side chains of different sizes synthesized by different enzymes, or are they produced by the same enzyme whose activity responds to environmental inputs? Finding plant mutants that lack a certain type of side chain or substituent will be crucial to answering these questions. The simple and well annotated genome of the model system Arabidopsis, as well as its valuable and easily accessible mutant collections can benefit us studying the roles of RG-I in plant growth and development. Studies of enzyme biochemistry are often restricted by the availability of enzyme substrates. We speculate that degrading the native RG-I isolated from different plant sources will provide a valuable substrate library for testing and characterizing enzymes that may be involved in RG-I biosynthesis.

## Experimental Procedures

### Chemicals and Reagents

All chemicals and reagents were purchased from Sigma (USA), Genessee Scientific (USA), or Fisher Scientific (USA) unless otherwise noted. All dialysis steps were carried out using Spectrum™ Spectra/Por™, 3500 Dalton MWCO tubing (Spectrum Chemical Mfg Corp, USA) unless otherwise indicated.

### Plant materials and growth conditions

*Arabidopsis thaliana* ecotype Columbia-0 (Col-0) was used in all experiments. Seeds were placed on a soil mixture consisting of a 1:1 ratio of Metromix 830 (Sungro, USA) and Vermiculite, which was supplemented with Plant Food (Sta-Green, USA) and Osmocote Smart-Release Plant Food Flower & Vegetable (Scotts, USA). Seeds were stratified for 2 days at 4°C before being moved to a walk-in growth chamber (Conviron, Canada). Plants were grown under an 18 hours light/ 6 hours dark cycle at 21°C with a light intensity of 120 µmol photons m^−2^ s^−1^.

Rosette leaves were collected from 4-week-old plants. Flowers (including fully opened flowers and flower buds), siliques, and inflorescence stems were collected from 6-week-old plants. Collected tissues were stored at -80°C before being processed.

### Preparation of alcohol insoluble residue (AIR) and isolation of pectins

Plant cell walls were prepared as alcohol insoluble residues (AIR) and fractionated according to established methods with some modifications (Barnes et al. 2021). Briefly, frozen plant tissue was homogenized in 80% ethanol using a POLYTRON® Homogenizer (Brinkmann Instrument). The homogenate was filtered with 50 μm nylon mesh. The residue was washed with 80% ethanol and then stirred in a 1:1 ratio of chloroform/ methanol overnight. The next day, residue was filtered with a 50 μm nylon mesh and washed first with 1:1 ratio of chloroform/ methanol and then with acetone. The washed residue was left in a chemical fume hood to air dry.

To remove starch, AIR (1 g) was resuspended in 100 ml of 0.1M sodium acetate (pH 5) containing 120 μl of Spirizyme Excel (Novozymes) and 600 μl of Liquozyme SC DS (Novozymes) and incubated for 24 hr at 50°C and shaking at 200 rpm. De-starched AIR was filtered using 50um nylon mesh, washed with 0.1M sodium acetate (pH 5), and then resuspended in 100 ml of 0.5% ammonium oxalate (pH 5). The oxalate suspension was left at room temperature with agitation at 200 rpm for 24 hrs. To obtain the pectin enriched, oxalate soluble fraction and the corresponding oxalate-treated AIR, the suspension was centrifuged at 2683g for 10 min. The supernatant was filtered with a GF/B glass microfiber filter (Whatman). The pellet was washed three times with 0.5% ammonium oxalate (pH 5). Each time, the pellet was resuspended in 50 ml of ammonium oxalate, centrifuged at 2683g for 10 minutes, and the resulting supernatant was filtered with a GF/B filter. The filtrates from filtering the supernatant of the initial oxalate suspension and from washing the pellet were combined. The combined filtrate and the oxalate-treated AIR residue were used in this study and termed the oxalate soluble fraction (OSF) and the oxalate insoluble residue (OIR), respectively.

### Preparation of RG-I from oxalate solubilized pectin

The volume of the oxalate soluble fraction was reduced to 70-120 ml using a rotary evaporator (Brinkmann) with the temperature of the water bath set at 37°C. The concentrated oxalate soluble fraction was dialyzed against deionized water for 2 days by using 3.5K MWCO dialysis tubing (Ward’s science) freeze-dried on a lyophilizer (VirTis). Oxalate soluble material (10 mg) was dissolved in 4 ml of 100 mM ammonium formate (pH 5.5) containing 4 μl endo-polygalacturonanase M2 (EPG, Megazyme). The reaction was conducted at 40°C with gentle rotation for 24 hrs. RG-I was isolated by separation of the resultant EPG labile pectin fragments by size exclusion chromatography (SEC). In brief, the EPG digest was filtered with a 0.45um nylon spin filter (ThermoFisher Scientific). SEC was carried out on a 1260 infinity II LC system (Agilent), which consists of a quaternary pump (G7111B, Agilent), a variable wavelength detector (G7114A, Agilent), a refractive index detector (G7162A, Agilent), and a fraction collector (G1364F, Agilent). A Superdex 75 Increase 10/300 GL column (Cytiva) was used with 50 mM ammonium formate (pH 5) as the mobile phase, and run isocratically at a flow rate of 0.4 ml/ min. Peaks corresponding to RG-I were collected and repeatedly freeze-dried and dissolved in Milli-Q water to remove ammonium formate.

### Preparation of RG-I from oxalate treated cell wall residues

Wall material that had been previously extracted by oxalate was resuspended in 90 ml of 100 mM ammonium formate, pH 5.5, containing endo-polygalacturonanase M2 (EPG, Megazyme; 12 μl) to release RG-I. The reaction was conducted at 40°C with gentle rotation for 24 hrs. The slurry was filtered with a 50 μm nylon mesh followed by a GF/B filter. The EPG treated residue underwent a second treatment with EPG under the same conditions. The filtrates were combined, concentrated using a rotary evaporator at 37°C, and then freeze-dried. The resulting dry material containing pectin fragments was dissolved in 6 ml Milli-Q water, centrifuged to remove insoluble material, and the supernatant was filtered using a 0.45 μm nylon spin filter. RG-I was separated by SEC and prepared for analysis following the same method as described above. Sugar linkage analysis and NMR analysis showed that the prepared RG-I still contained a significant amount of HG. To obtain pure RG-I, we digested the initial RG-I again with a combination of EPG and pectin methylesterase (PME; gift of Hans Peter Heldt-Hansen, Nova Nordisk A.S, Bagsvaerd, Denmark). The product was again separated via SEC.

### Expression and purification of BT4175 (RG-I lyase), BT4668 (endo-β1,4-galactanase), and BT0368 (endo-α-1,5-arabinanase)

Plasmids that contain BT4175, BT4668, and BT0368 genes were kind gifts from Dr. Harry J. Gilbert (Luis et al. 2018). *Escherichia coli* (*E.coli*) strain TUNER was transformed with the plasmid and grown to mid-exponential phase (OD600: 0.6-0.8) before induction with 0.1 mM isopropyl β-D-galactopyranoside (IPTG). Protein induction was carried out overnight at 16°C with shaking at 160 rpm. Recombinant proteins were purified using HisPurTM cobalt resin (ThermoFisher Scientific). BT4175 was buffer-exchanged, using dialysis, into 50 mM HEPES-Na, 400 mM NaCl, 5 mM CaCl_2_, pH 7. BT4668 and BT0368 were buffer-exchanged, using dialysis, into 20 mM MES, 150 mM NaCl, pH 6.5. Following dialysis, BT4175 and BT0368 were concentrated to 2mg/ml using Amicon Ultra centrifugal concentrators (MWCO: 30kDa, EMD Millipore), while BT4668 was concentrated to 2mg/ml using a concentrator with MWCO of 10kDa. Protein concentration was measured by Bradford assay (Bio-Rad).

### Enzymatic assays

BT4175 is an RG-I lyase that has been previously shown to accommodate glycans appended to backbone rhamnose units (Luis et al. 2018). To evaluate the activity of BT4175, two RG-I substrates were used: Arabidopsis seed mucilage and potato RG-I. Non-adherent Arabidopsis seed mucilage is reported to be unbranched RG-I. RG-I produced from potato pectic fiber (Megazyme) is a branched RG-I containing short galactan side chains. RG-I (10mg/ml) substrate was prepared in Milli-Q water. Reactions contained BT4175 (1.6 ug) and 100 ug substrate. BT4175 is an RG-I lyase that cleaves the backbone generating an unsaturated double bond between the C4 and C5 of the nonreducing end GalA residue. The formation of double bonds during digestion can be monitored by UV absorbance at 235nm (Kofod et al. 1994), which was done by using a Take3 plate (Agilent BioTek) and an Epoch microplate spectrophotometer (Agilent BioTek).

The pH optimum was determined using Arabidopsis seed mucilage as the substrate. Buffers were as follows: 75 mM sodium acetate, pH 5; 75 mM MES, pH 6; 75 mM HEPES sodium, pH 7; 75 mM HEPES sodium salt, pH 8; and 75 mM Tris HCl, pH 9. To evaluate the divalent cation dependency of BT4175, the enzyme was first incubated in 5 mM EDTA on ice for one hour. Subsequently, reactions were set up in 75 mM HEPES sodium salt, pH 7 as the reaction buffer and mucilage as the substrate. MnCl_2_, MgCl_2_, or CaCl_2_ were evaluated individually and added to the reactions at a final concentration of 10 mM. The effect of temperature on BT4175 lyase activity was measured at 4°C, 24°C, 37°C, and 42°C. 75 mM HEPES sodium salt, 10 mM MnCl_2_, pH 7 and Arabidopsis mucilage were used as the reaction buffer and substrate, respectively. Product formation was monitored after 30 minutes, 1 hour, and overnight.

To fragment RG-I isolated from Arabidopsis oxalate insoluble residue (OIR), HEPES sodium salt, pH 7, containing 10 mM MnCl_2_ was selected as the reaction buffer. 20 mM HEPES sodium salt buffer was used as the reaction buffer. Reactions were conducted at 37°C overnight. 6 ug of BT4175 was used to digest 150 ug of RG-I. Chemically de-esterified RG-I was also prepared for analysis. To achieve this, RG-I (1 mg) was dissolved in 100 μl of Milli-Q water. Concentrated NaOH solution was carefully added to the dissolved RG-I while stirring until pH reached 12.3-12.7. Chemical de-esterification was conducted on ice for one hour. Subsequently, the pH of each RG-I sample was adjusted close to its original pH by using HCl.

BT4668 is an endo-galactanase (Luis et al. 2018). To test the activity of BT4668, potato galactan (Megazyme) was used as the substrate, which was prepared in Milli-Q water as a 10 mg/ml stock solution. BT4668 (2 μg) was used to digest 30 μg of substrate in 20 mM sodium acetate, pH 5. Reactions were conducted at 37°C overnight. The reaction products were analyzed by MALDI-TOF MS (smartfleX, Bruker). The same conditions and method were used to analyze the digests of RG-I isolated from Arabidopsis oxalate insoluble residue (OIR).

BT0368 is thought to cleave the α-1,5-arabinofuranosyl residues of the pectic arabinan (Luis et al. 2018). To evaluate the activity of BT0368, sugar beet arabinan (Megazyme) was used as the substrate, which was made in Milli-Q water as a 10mg/ml stock. BT0368 (2.7 μg) was used to digest 62 μg substrate. The reaction buffer was 20 mM MES, pH 6. Reactions were conducted at 37°C overnight. Release of arabinose was detected by high-performance anion-exchange chromatography with pulsed amperometric detection (HPAEC-PAD) on a Dionex ICS-6000 system (ThermoFisher Scientific), as described (Pellerin et al. 1996). Briefly, the digests were filtered through 0.45 μm nylon spin filters (Thermo Fisher Scientific), and analyzed using a Dionex CarboPac PA1 column (ThermoFisher Scientific; 4x250mm; 10um). The column was eluted at 1ml/min with 32 mM NaOH (0-15min) followed by a gradient of sodium acetate from 0 to 0.25 M in 100 mM NaOH (15-35 min), a gradient of sodium acetate from 0.25 to 1M in 100 mM NaOH (35-45 min), and then with 1 M sodium acetate in 100 mM NaOH (45-48 min). The column was equilibrated into 32 mM NaOH within 3 min and then further equilibrated for 9 min prior to the next injection. The same conditions and method were used to analyze the digests of RG-I isolated from Arabidopsis oxalate residue.

### Linkage analysis

Glycosyl linkage analysis was performed by combined gas chromatography-mass spectrometry (GC-MS) of the partially methylated alditol acetates (PMAAs) derivatives produced form the samples. The procedure was described in detail by Black et al. (Black et al. 2023). Briefly, the samples were acetylated using N-methylimidazole and acetic anhydride in the ionic liquid 1-ethyl-3-methylimidazolium acetate ([Emim][Ac]). The acetylated samples were dialyzed against DI water for 2 days in a 3-kDa MWCO dialysis membrane. The samples were then methylated using potassium methylsulfinylmethylide (dimsyl potassium) base. Following extraction with dichloromethane (DCM), the carboxylic acid methyl esters were reduced using lithium aluminum deuteride (LiAlD4) in tetrahydrofyran (THF). After a second dialysis as before, the samples were remethylated using two rounds of treatment with sodium hydroxide and methyl iodide. The samples were then hydrolyzed using 2 M trifluoroacetic acid (TFA), reduced with sodium borodeuteride (NaBD_4_), and acetylated using acetic anhydride/ TFA. The resulting PMAAs were analyzed on an Agilent 7890A GC interfaced to a 5975C MSD (mass selective detector, electron impact ionization mode); separation was performed on a 30-m Supelco SP-2331 bonded phase fused silica capillary column.

### NMR spectroscopy

Samples for NMR spectroscopy were prepared by dissolving pectin (2–3 mg) in D_2_O (99.9% D, Sigma) with 60 nmol DSS-d_6_ (99% D, CIL) and lyophilizing. Each sample was then dissolved in 42 or 510 μL D_2_O (99.96% D, CIL) and transferred into a 1.7 mm (800 MHz) or 5 mm (600 MHz) NMR tube, respectively. NMR data were acquired at 30 °C on a Bruker NEO spectrometer with a 1.7 mm TCI cryoprobe (1H, 799.71 MHz) or a Bruker Avance III with a 5 mm TCI cryoprobe (1H, 600.13 MHz) as indicated. Standard pulse programs included in the spectrometer library were used to acquire 1D ^1^H and 2D ^1^H,^1^H-COSY, ^1^H,^1^H-TOCSY, ^1^H,^1^H-NOESY, ^1^H,^1^H-ROESY, ^1^H,^13^C-HSQC, ^1^H,^13^C-HMBC and ^1^H,^13^C-HSQC-TOCSY experiments. Quantitative ^1^H NMR data were collected with a total recycling delay of 60 s. Mixing time was set to 80–120 ms for NOESY and to 200 ms for ROESY experiments. Spin-lock time was 70 and 120 ms for TOCSY, and 80 ms for HSQC-TOCSY experiments. NMR data were processed in Topspin 4.0.5 (Bruker), analyzed using CCPN Analysis 2.4 (Vranken2005), and rendered for figures in MestReNova 14.2.3 (Mesterelab Research). ^1^H and ^13^C chemical shifts were referenced to the respective DSS signals at 0.00 ppm.

### Liquid chromatography with tandem mass spectrometry (LC-MS/MS) analysis

RG-I oligosaccharides generated from digesting Arabidopsis seed mucilage and potato RG-I by polysaccharide lyase BT4175 were analyzed by LC-MS/MS using a Vanquish UHPLC system (ThermoFisher Scientific) connected to a QExactive orbitrap mass spectrometer (ThermoFisher Scientific). Briefly, separation was carried out on a Supel Carbon LC (Supelco; 2.1x100 mm; 2.7 µm) with separation conditions as follows at 0.3ml/min: 0-3 minutes the buffer was held at 90% acetonitrile in 0.1% formic acid, dropped to 50% acetonitrile and held for 8min, and increased from 50% acetonitrile to 90% acetonitrile for another 12min to equilibrate the column for the next run. The first 10 min of the separation was fed into the mass spectrometer for analysis while the equilibration was diverted to waste. A data-dependent program was used for acquisition where the precursor ion scan was acquired at 120k resolution followed by top-down fragmentation of high-to-low intensity m/zs by stepped HCD at 30k resolution. For C18 separations, a commercial C18 column (Zorbax Eclipse XDB-C18; 2.1x150 mm; 1.8 µm) was used with a linear gradient of 1% to 99% acetonitrile in 0.1% formic acid over 22 min. The column was cleaned for 2 min with 99% acetonitrile followed by 4 min of reconditioning using 1% acetonitrile. Data was processed using Freestyle software (ThermoFisher Scientific). Prior to MS analysis, digests of mucilage and potato RG-I by BT4175 were filtered through 0.45 μm nylon spin filters.

### Matrix-assisted laser desorption/ionization time-of-flight mass spectrometry (MALDI-TOF MS) analysis

Proteins and salt in the digests of RG-I isolated from Arabidopsis oxalate insoluble residue (OIR) by BT4175 were removed using a Dionex CarboPac PA20 column (ThermoFisher Scientific; 4x250 mm; 10 μm) using a 1260 infinity II LC system (Agilent). Briefly, digests were filtered through 0.45 μm nylon spin filters (ThermoFisher Scientific) and then manually injected onto the PA20 column. The separation conditions were as follows at 0.5 ml/min: a Milli-Q water wash for the first 10 min, followed by elution with 0.5 M ammonium formate at pH 5 for 20 min, then the column was equilibrated in Milli-Q water for 20 min. RG-I oligos were collected based on UV absorbance at 245 nm. The collected RG-I oligos were repeatedly lyophilized and dissolved in Milli-Q water to remove ammonium formate. Purified RG-I oligos were finally dissolved in 20 μl Milli-Q water and analyzed by MALDI-TOF MS.A solution of RG-I oligos (1 μl) was diluted 10-fold with Milli-Q water. Nafion 117 (0.1 μl; Sigma-Aldrich, USA) was applied onto a MTP 384 target plate ground steel (Bruker) and air-dried first. Then 1 μl of the diluted digest was mixed on the plate with 1 μl of DHB matrix (20mg/mL 2,5-dihydroxybenzoic acid (DHB) made in 50% methanol). Samples were dried using a hair dryer (Conair). Analyses were carried out on a RapifleX system (Brucker) in positive ion mode. Data was processed in flexAnalysis (Brucker).

## Author Contributions and Acknowledgments

L.Z., J.V., I.M.B., and S.A.H performed the experiments. L.Z., J.V., I.M.B., S.A.H and C.H. designed experiments, interpreted and analyzed data, created figures and tables and and wrote the paper. P.A. conceived the study, acquired funding, supervised work, and edited the manuscript with input from all authors. B.R.U. conceived the study, acquired funding, designed experiments, interpreted data, supervised work, and wrote and edited the manuscript with input from all authors.

## Supplementary data

Supplementary Figure S1. Glycosyl linkage analysis of oxalate soluble fraction (OSF) and RG-I made from oxalate insoluble residue (OIR).

Supplementary Figure S2. NMR spectra of the oxalate soluble fractions (OSF).

Supplementary Figure S3. Separation of RG-I made from oxalate soluble fraction (OSF) and oxalate insoluble residue (OIR) of Arabidopsis silique by size exclusion chromatography.

Supplementary Figure S4. NMR spectra of RG-I obtained from the oxalate insoluble residue (OIR).

Supplementary Figure S5. Regions of 2D TOCSY (70 ms) and NOESY (60 ms) spectra with correlation signals used to identify 3-*O* acetylation of Rha residues.

Supplementary Figure S6. HPAEC-PAD analysis of arabinose released by BT0368.

Supplementary Figure S7. MALDI-TOF MS analysis of galactobiose released by BT4668.

Supplementary Figure S8. BT4175 exhibited RG-I lyase activity on both unbranched RGI and branched RG-I.

Supplementary Figure S9. The effects of pH, cation, temperature, and incubation time on the activity of BT4175.

Supplementary Figure S10. Separation of RG-I oligosaccharides generated by BT4175 from potato RG-I by size exclusion chromatography.

Supplementary table S1. Chemical shift assignments of carbohydrate residues in Arabidopsis silique RG-I sample at pD 4.7 and 1.7 (select residues)

Supplementary table S2. Oligosaccharide structures generated by digestion with RG-I lyase BT4175 observed by MALDI-TOF MS.

## Funding

Isolation and analysis of RG-I performed in this work was supported by the U.S. Department of Energy, Office of Science, Basic Energy Sciences, grant number DE-SC0015662 to the Complex Carbohydrate Research Center. Research related to the generation of oligosaccharide substrates for functional characterization of glycoenzymes was supported by the U.S. Department of Energy, Office of Science, Biological and Environmental Research, Genomic Science Program grant no. DE-SC0023223 to B.R.U.

